# Game of Theories: An Agent-Based Model of How Variation in Contest Cost-Benefits Determines the Evolution of Contest Resolution Rules

**DOI:** 10.1101/2021.12.06.471375

**Authors:** Nelson Silva Pinto

## Abstract

Game-theory based models are used to understand rules that animals use to settle contests over indivisible resources. However, the empirical literature of contests indicates controversial support to models, with some species supporting different models and other species showing no support to any model. Since strategies used to resolve contests may have different associated costs, it is possible that different conditions have determined the evolution of distinct assessment strategies used by animals. We used an agent-based model to explore the importance of the following conditions: resource availability, probability of reproduction with resource, and damage costs on evolution of assessment strategies. We used self- and mutual-assessment models as a heurist framework to build agents with different assessment strategies. In our model, agents competed for resources in scenarios with different combinations of resource availability, probability of reproduction with resource, and damage costs. We found that agents following the self-assessment with damage strategy were prevalent in scenarios with no probability of reproduction without the resource, independently of other variables. We also found that agents following the non-aggressive strategy occurred in all scenarios. However, agents using the non-aggressive strategy were prevalent only in scenarios with probability of reproduction with the resource. Finally, we observed that agents using mutual-assessment occurred only in a scenario with high risk of damage, low availability of resources, and with probability of reproduction without the resource. These results indicate that agents following the self-assessment with damage and non-aggressive strategies may be able to stay at most scenarios.

## INTRODUCTION

Models based on game theory represent one of the most prolific theoretical frameworks used to understand how individuals behave during contests and how they determine the winner of the interaction (Hardy & Briffa 2013). Some of these models predict the existence of contests in which rivals are unable to impose direct costs to each other (Mesterton-Gibbons *et al*. 1996). In such contests, the giving up process occur when one of the rivals reaches an internal cost threshold of time and/or energy (self-assessment) and flee from the contest. Other models also predict the existence of an individual cost threshold to quit the interaction. However, the individual cost threshold may be influenced by costs imposed by rivals during the contest (cumulative assessment, Payne 1998). There are also models that predict that animals can evaluate the fighting ability (RHP-resource holding potential, Parker 1974) of rivals and compare it to its own RHP (mutual-assessment, Enquist & Leimar 1983). According to mutual-assessment, as soon the individual perceives itself as the weaker rival, it may flee from the contest. Finally, there are other recent models (e.g. Mesterton-Gibbons & Heap 2014) that propose a process in which individuals may use a combination of self- and mutual-assessment during different phases of the contest.

Empirical studies testing the explicative power of models have found support for different models in different species (e.g. Palaoro *et al*. 2014; Guillermo-Ferreira *et al*. 2015) or patterns not predicted by any model (e.g. Rillich 2007). However, there is no explanation about the possible causes of such variation in contest decision rules among species. It is possible that variations in the conditions in which the species had evolved have favoured different strategies. For example, we can expect that animals fight harder in regions with lower abundance of the resource. We also can expect that in such conditions, animals evolve to use weapons or to provoke injuries in rivals during contests. Then, it is important to investigate if it is possible that different species that evolved in similar environmental and social conditions present convergent evolution of strategies of contest resolution.

In contests based on displays without physical damage, only energetic or time costs are expected (Mesterton-Gibbons *et al*. 1996). Contests in which animals can inflict costs on each other may present additional costs of damage or death (Payne 1998; Payne & Pagel 1996). Therefore, the costs in contests with damage infliction may be greater than those related to contests with displays. However, in both these situations, individuals are expected to pay costs until reaches its individual cost threshold. On the other hand, individuals using mutual-assessment strategies can evict the costs of staying in a contest that they may lose (Enquist & Leimar 1983). Therefore, the mutual assessment strategy can allow the individual to save energy that can be expended in more profitable contests. In such cases, mutual-assessment may represent a way to minimize costs. However, animals using mutual assessment during contests may experience greater costs than animals involved in contests with displays only. Thus, it is possible that the mean costs of mutual-assessment are greater than those observed for displays and smaller than those observed for damage infliction. For many strategies used during the life of an organism, it is expected that costlier behaviours may be favored when the associated profits are higher (e.g. Palminteri *et al*. 2016). Similarly, costlier contest behaviours may be favored in scenarios in which the profits are higher.

Animals get involved in contests over different resources, such as shelter, territories, food or mates. However, a great number of studies focused on contests over mating territories (e.g. Guillermo-Ferreira *et al*. 2015; Peixoto & Benson 2011; Prenter *et al*. 2006). In these contests, it is possible that the number of available territories, the willingness of the females to reproduce with non-owner males and damage costs may affect the costs/benefits balance of contests (Hardy & Briffa 2013). In situations in which there is a low number of defendable territories, one may expect that males fighting with damage infliction may be favored due to the ability to impose damage costs to rivals. On the other hand, it is possible that the costs associated with the contest are lower in situations in which females may sometimes reproduce with non-owner males and there is a great number of available territories. Then, males with different strategies may be favored, since they can reproduce even without the possession of a territory. The damage costs can also affect the cost/benefit balance during contests (Briffa & Elwood 2009). Individuals of different species possessing different abilities to cause injuries invest less in contests when the risk of damage is high, even when profits are high. Then, it is possible that due to different damage costs, animals use different rules of assessment.

Our objective was to evaluate how the number of available resources (that can be though as suitable areas to establish as a territory), the chance of reproduction with and without the resource possession and damage costs can favour the evolution of different assessment strategies in contests over territories. To accomplish our aims, we used simulations with a computational agent-based model (ABM, Grimm & Railsback 2005). In an ABM it is possible to model individual characteristics of organisms, then allowing to evaluate how differences in individual traits can influence the interactions. In our ABM the agents differ in both their RHP and assessment strategy attributes. We used the game-theory models (e.g. Enquist & Leimar 1983; Mesterton-Gibbons *et al*. 1996; Payne 1998) as the basis to build different assessment strategies used by agents during contests. To build our set of strategies, we decomposed the assumptions of the models into dicothomic variables. The decomposition allowed us to buid a set of nine different combinations of assessment strategies used by our agents during contests (Table 1). Then, we proposed the following hypothesis. Contests solved according to self-assessment and damage infliction may be favored in scenarios with lower availability of resources and without probability of reproduction without the resource. Contests solved according to mutual-assessment may be favored in scenarios with intermediate availability of resources and with probability of reproduction without the resource. Contests solved according to self-assessment without damage infliction may be favored in scenarios with high availability of resources and with probability of reproduction without the resource.

**Table 1.** Summary of possible assessment strategies that agents could use in our model. Here we used different combinations of self-assessment, mutual-assessment and ability to inflict damage to produce agents following nine different assessment strategies (see strategy description). The value 0 indicate the absence of the characteristic in a given strategy and 1 refers to presence of the characteristic. Agents using the desperado without damage strategy do not perform any assessment and fight until death. Agents using desperado with damage strategy do not perform any assessment and fight until death but can cause damage to other agents. Agents using the self-assessment without damage do not inflict damage and fight until reaching the cost threshold. Agents using the mutual-assessment without damage can perform mutual assessment and fight until perceiving itself as the weaker rival. Agents using the mutual-assessment with cost threshold can perform mutual assessment, however the agent gives up if it reaches its threshold before concluding mutual-assessment. Agents using the self-assessment with damage fight until reaching a costs threshold and they are able to inflict damage. Agents using the mutual-assessment with damage can perform mutual assessment and also can inflict damage. Agents using the mutual-assessment with cost threshold and damage strategy can perform mutual assessment, however the agent give up if it reaches its threshold before concluding mutual-assessment, they also can inflict damage. Agents using the non-aggressive strategy never fight and leave the resource when confronted by another agent.

| Ability to inflict damage | Mutual assessment | Self assessment | Strategy name |
| --- | --- | --- | --- |
| 0 | 0 | 0 | desperado without damage |
| 0 | 0 | 1 | self-assessment without damage |
| 0 | 1 | 0 | mutual-assessment without damage |
| 0 | 1 | 1 | mutual-assessment with cost threshold |
| 1 | 0 | 0 | desperado with damage |
| 1 | 0 | 1 | self-assessment with damage |
| 1 | 1 | 0 | mutual-assessment with damage |
| 1 | 1 | 1 | mutual-assessment with cost threshold and damage |
| - | - | - | non-aggressive |

## METHODS

### 1. Model description

We build an ABM to investigate the effect of different combinations of damage costs, availability of resources and probability of reproduction with and without resource possession on evolution of different assessment rules. Our ABM simulated populations of *N* agents varying in RHP and assessment strategies, living in an environment with limited resources for which the agents competed aggressively. Our model incorporated rules of contest resolution that are based on RHP-correlated asymmetries. Uncorrelated asymmetries (e.g. respect for ownership, Sherratt & Mesterton-Gibbons 2015; Mesterton-Gibbons & Sherratt 2016) may be important to contest resolution in some species (Kemp & Wiklund 2004). Nonetheless, even in contests in which rivals use uncorrelated asymmetries, RHP-correlated asymmetries still play a role (Kasumovic *et al*. 2011). Therefore, the parameters and rules of assessment used in our model allowed to adequately explore the rules for contests resolution.

We used the game-theoretic models (Enquist & Leimar 1983; Mesterton-Gibbons *et al*. 1996; Payne 1998) as the basis for the definition of all strategies of contest resolution that the agents may present in the model. These models were created as alternative explanations for animal assessment strategies. Therefore, for the modeling of alternative strategies of assessment to make sense, we decomposed the strategies described by the models in three binary traits: self-assessment, opponent-assessment, and damage infliction (Table 1). Then, we used these three binary traits to build the setup of assessment strategies that defined assessment rules that the agents used during contests. By decomposing the strategies in simple binary traits, we were able to develop a unified model in which such interactions were possible (see details below).

Each simulation referred to one season of one population containing 1800 agents. The number was partitioned to provide the same initial proportion of agents with different assessment strategies (200 agents for each strategy). The offspring inherited the assessment strategy of the parental agent. The agents varied in its RHP. The RHP was the variable related to the fighting ability. As the agents fought and the time passed, the decreasing in the RHP variable and costs threshold was updated. The RHP also was related to investment in reproduction and to the life of the agent. As the value of the RHP approached zero, the value that an agent could invest in reproduction decreased. When the value reached zero, the agent died.

The RHP for each agent was drawn from a truncated normal distribution with mean(M)=100±standard deviation(SD)=5 and minimum value=0. The agents also varied in their cost threshold, that was related to RHP. Our model was focused on contests for the possession of territories. However, the model can be applied to contests over other resources. For example, in contests over shelters it would be possible to evaluate effects of shelter availability and how the shelter possession may affect the survival of the individuals. Resources did not vary in quality among themselves, although they varied in availability (Table 2).

**Table 2.** Number of resources, damage costs per round of contest and probability of reproduction with and without the resource possession. Each scenario had three combinations of parameters.

| Scenario | Number of resources | Damage costs infliction | Probability of reproduction |  |
| --- | --- | --- | --- | --- |
|  |  |  | Without resource | With resource |
| 1 | 180 | 0.1 | 0 | 0.9 |
|  | 450 | 0.1 | 0 | 0.9 |
|  | 900 | 0.1 | 0 | 0.9 |
| 2 | 180 | 0.5 | 0 | 0.9 |
|  | 450 | 0.5 | 0 | 0.9 |
|  | 900 | 0.5 | 0 | 0.9 |
| 3 | 180 | 0.1 | 0.3 | 0.9 |
|  | 450 | 0.1 | 0.3 | 0.9 |
|  | 900 | 0.1 | 0.3 | 0.9 |
| 4 | 180 | 0.5 | 0.3 | 0.9 |
|  | 450 | 0.5 | 0.3 | 0.9 |
|  | 900 | 0.5 | 0.3 | 0.9 |

### 2. Process overview

The model was not spatially explicit and time was divided into discrete time steps (Fig. 1). There were three nested time scales in the model. The finest scale was the contest time-scale, the middle time scale was the ecological one, and the broadest was the generational time scale. Each contest was composed of several steps in the contest time scale, so that one step in this scale may represent few seconds to minutes. A time-step in the ecological scale represents the time needed for an individual to locate a resource and try to settle within it. Finally, a time-step at the generational time scale represents a whole generation: a breeding season plus the time needed for the offspring to reach reproductive maturity.

**Figure 1.**
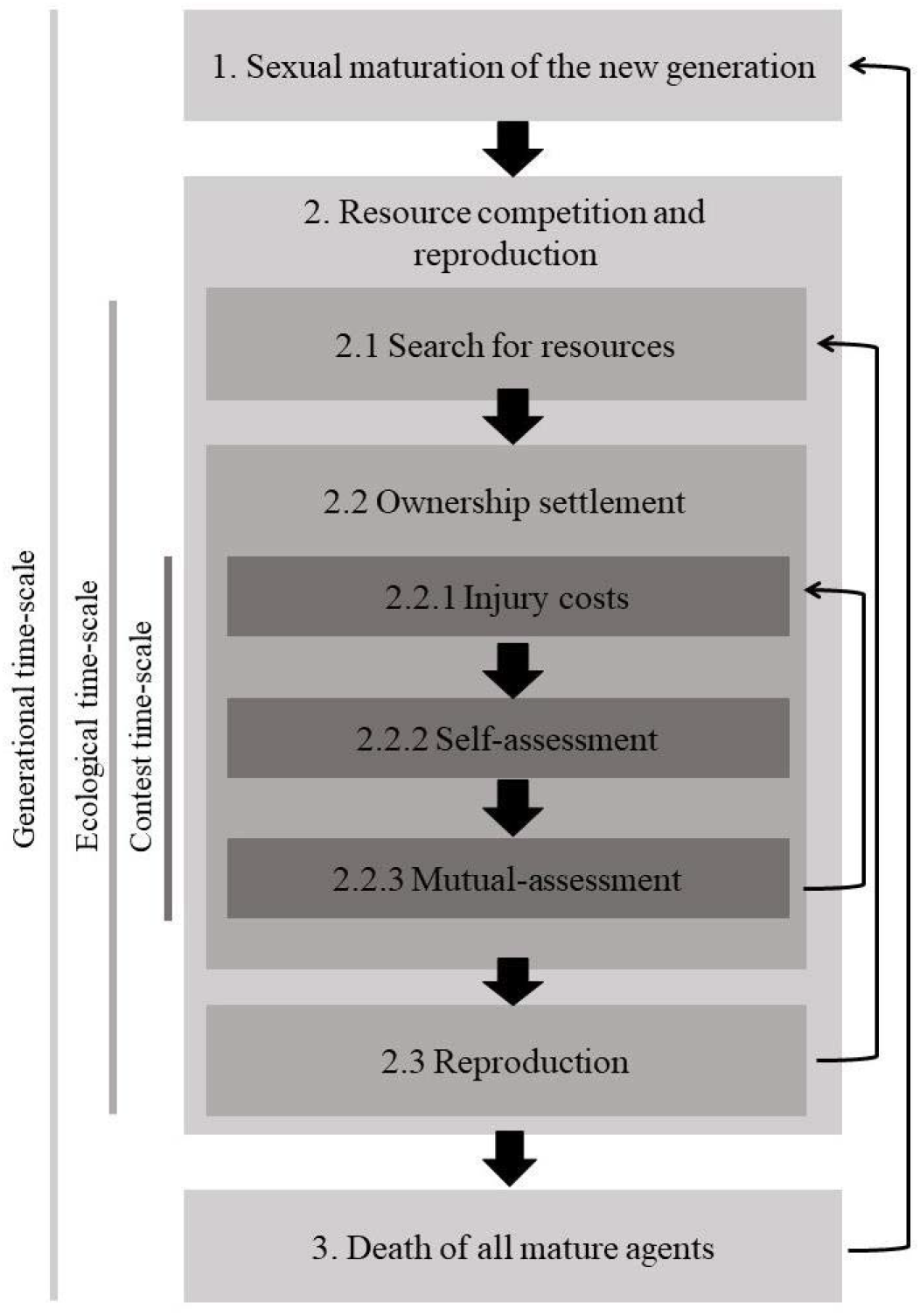
Overview of simulation used in our model. Shades of grey indicate each phase and the time-scale in the simulation. The darker shade of grey indicates the contest time-scale. At this phase, agents contest each other using inherited assessment rules to stablish the ownership over resources. The middle-dark shade of grey indicates the ecological time-scale. At this phase, all mature agents may search for resources. All contests occur nested at the ecological time-scale. At the ecological time-scale, agents also may reproduce. The lighter shade of grey represents the generational time-scale. At this phase, maturation of new generations occurs. All phases of the contest time-scale and ecological time-scale occur nested in the generational time-scale. Broad arrows indicate the flowchart of phases in the model. Thin arrows indicate the iterated process at each time-scale. At the end of the simulation, all remaining mature agents die.

At the beginning of each simulation, there was a probability of each agent finding a resource. If the agent found a resource at the first time-step, it became the owner. If two agents found a resource at the same time, they disputed the resource. After this, for all agents that had not found resources, there was a probability to find a resource, representing the search for resources. If the agent found an unoccupied resource, then it became the new owner. On the other hand, contests occurred if the agent found an occupied resource. After the contest resolution, the winner stayed with the resource and the loser tried to find a new resource.

#### 2.1. Contest time scale

Whenever an agent found a resource already being defended by another agent, a contest occurred. Contests were composed by a series of rounds, each lasting one time-step in the contest time scale. The whole duration of the contest was defined by the contestants’ strategies and RHP values (see below). At each round, the following phases occurred: (1) costs of staying in the contest and damage infliction were calculated, (2) self-assessment and (3) mutual-assessment (Fig. 1). During phase 1, the agents payed the costs of staying in the contest and the costs of received damage per round of the contest. These costs were updated at each round of the contest. As the costs were accumulated at each round of the contest, these costs were subtracted from the RHP variable, then the RHP were updated at each round of contest. At each round the costs were compared to the cost threshold of the individuals. Individuals gave up when the accumulated costs were equal or greater than the cost threshold.

During phase 1, both contestants also had the opportunity to inflict damage in each other if they had this capability. The damage was established as low (0.1) and high (0.5). The agent able to inflict damage paid a cost related to possessing the ability to inflict damage. An individual with a self-assessment trait equal to one (Table 1) was able to assess how much *RHP* it had spent in this contest, if the value was above its threshold *t*, the agent would give up. The variable *t* was proportional to the RHP of the agent. An agent with self-assessment 0 (Table 1) never gave up in this phase. This agent could invest all the RHP in a contest and die. During the mutual-assessment, agents with this ability could perform an estimate of the opponent’s RHP. The estimate of the opponent’s RHP had an associated error (Enquist & Leimar 1983). Thereafter, agents evaluated if the rival was stronger. If the agent judged that the rival was stronger, it abandoned the contest. Otherwise, the agent stayed in the contest.

In our model, we first calculated the estimate of the RHP for each agent in the contest. First, we used an equation for agents with mutual-assessment trait to evaluate the RHP of rivals with an error. The error variable was a characteristic of the agent. Then, each agent who could perform mutual-assessment had an error associated with the assessment of the RHP of the rivals. The value of the error (*E*) was obtained from a normal distribution with mean 0 and standard deviation 5. We used the error value calculated for each agent in equation 1. At all rounds of the contest, the agent with mutual-assessment ability performed estimates of the RHP of the rival. The closer to zero the value of the error was, the more accurately the agent could evaluate the rival’s RHP. Then, equation 1 represents one round of assessment that the agent (i) made of the RHP of the rival (j), added to the error of evaluation for the agent (i).

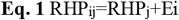

After the evaluation of the rival’s RHP, the agent performed the relative RHP assessment. The relative RHP assessment was simulated as follows. First, the agent compared the RHP of the rival with its own RHP. Then, there was the calculation of the measure of relative RHP (rij, equation 2). Equation 2 represents one round of calculation of the relative RHP for the agent *i* in relation to the agent *j*.

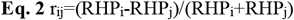

In the next phase the agent should evaluate if the rival agent was stronger than itself. Then, to perform this evaluation, first we calculated a probability of estimating the rival as the stronger one. The equation to calculate this probability used the relative assessment value for the agent *i*. Equation 3 was used to estimate the probability of performing a correct assessment of the rival, for a given agent *i*. If the value obtained in equation 3 was equal or greater than the limiar value, then the agent evaluated the rival as the greater one and it would flee from the contest. On the other hand, if the value obtained in the equation 3 was lower than the limiar value, then the agent would persist in the contest, where relative RHP for the agent *i* in relation to the agent *j*.

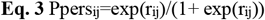

#### 2.2. Ecological time scale

At the ecological time scale, the agents searched for resources. After the search, it is possible that they found resources and evaluated if the resource had an owner (Fig. 1). The probability to find a resource was 0.5. After all contests for all agents were solved, the agents could reproduce (Fig. 1). This procedure allowed us to obtain the remaining energy that a given agent can expend in reproduction and also allowed us to verify if there were some agent with RHP=0. It is important to note that the reduction and the updating in the values of the RHP for agents affected the agent contest threshold. If one agent reached RHP=0, then it was assigned as dead and was excluded from the following steps. At each ecological time-step, agents paid a common cost (Cm). This cost referred to the energy expenditures of being alive during a round of the ecological time scale. The Cm parameter is fixed to all agents and generations as Cm=0.25. After the removal of all died agents, the agents proceeded to reproduction. There were different probabilities of reproduction (Table 2). Agents possessing the resource had 90% of chance of reproduction. Agents without the resource possession had either 0% or 30% of chance of reproduction. After reproduction, all agents died.

#### 2.3 Generational time scale

During the generational time scale hatching and maturation of the new generation took place (Fig. 1). All new agents were selected randomly from the offspring produced by parental agents. The random selection allowed agents with the most successful strategies to be better represented at the beginning of each new generation. It is important to note that the offspring inherited the assessment rule from its parental agent without any mutation. After the hatching of the new generation, the agents could invest in all phases at ecological time scale and contest time scale. Finally, after the agents had accomplished all tasks in the model, all mature agents died and the new generation took place.

### 3. Initialization and analyses of simulation data

At the beginning of simulations, we assigned the values to model parameters according to Table 3. There are four scenarios, each one with three possible combinations of number of resources, damage costs per round of contest and probabilities of reproduction. We defined resources availability as: low (180), intermediate (450) and high (900) number of resources. The number of resources can influence the number of contests that an agent experience during its lifetime. The damage cost per round of contest had two values explored in our model: low (0.1) and high (0.5). We used two different values of damage costs to explore the importance of costs that individuals can impose in each other during contests. Then, damage refers to the ability to affect the cost threshold of the rival, increasing the rate in which the rival reaches its cost threshold (Payne 1998).

**Table 3.** List of all acronyms used in the model as variables for all simulations and the description of each variable. We also present all values for the variables used in our simulations.

| Parameter | Description | Values used in simulations |
| --- | --- | --- |
| N | Number of agents per simulation | 1800 |
| G | Number of generations | 150 |
| R | Number of resources per simulation | 180;450;900 |
| D | Damage per round of contest | 0.1;0.5 |
| S | Error in opponent assessment | 5 |
| H | Intercept of the persistence relation in mutual assessment equation | 1.5 |
| J | Effect size of the persistence relation in mutual assessment equation | 20 |
| Cf | Cost to stay one round of the contest | 0.025 |
| Cm | Cost of being alive | 0.25 |
| p_resource | Probability of reproduction with resource | 0.9 |
| p_poor | Probability of reproduction without resource | 0;0.3 |
| Csa | Cost to perform self-assessment | 1.5 |
| Cma | Cost to perform mutual-assessment | 1.5 |
| Cinj | Cost to possess ability to inflict damage | 1.5 |
| F | Probability of an agent finding a resource at any time | 0.05 |
| Qmean | Mean of RHP for agents | 100 |
| Qsd | Standard deviation of RHP for agents | 5 |
| TRmean | Mean of costs threshold for agents | 0.2 |
| TRsd | Standard deviation of threshold for agents | 0.2 |

We used one value for the probability of reproduction with the resource possession (P= 0.9). For the probability of reproduction without the resource possession we used two different values (P=0 and P=0.3). We used these values to simulate situations in which animals can only reproduce in possession of resources or for which there are some chance of reproduction without the possession of resource. Our simulation consisted of 150 generations. We implemented the model in R software version 3.2.5 (R Development Core Team, 2011). All codes used in simulations were presented in the supplementary material.

## RESULTS

### **First scenario:** low damage costs per round of contest (D=0.1), no probability of reproduction without resource possession

For low resource availability (R=180), 92.44% of the agents presented self-assessment with damage strategy (Fig. 2a), 5.23% presented the non-aggressive strategy (Fig. 2a) and 2.33% presented the desperado with damage strategy (Fig. 2a). For intermediate resource availability (R=450), 81.56% of the agents presented self-assessment with damage strategy (Fig. 2b), 9.83% presented the non-aggressive strategy (Fig. 2b) and 8.61% presented the desperado with ability to inflict damage strategy (Fig. 2b). For high resource availability (R=900), 75.22% of the agents presented self-assessment with damage strategy (Fig. 2c), 17.56% presented the desperado with damage strategy (Fig. 2c) and 7.72% presented the non-aggressive strategy (Fig. 2c).

**Figure 2.**
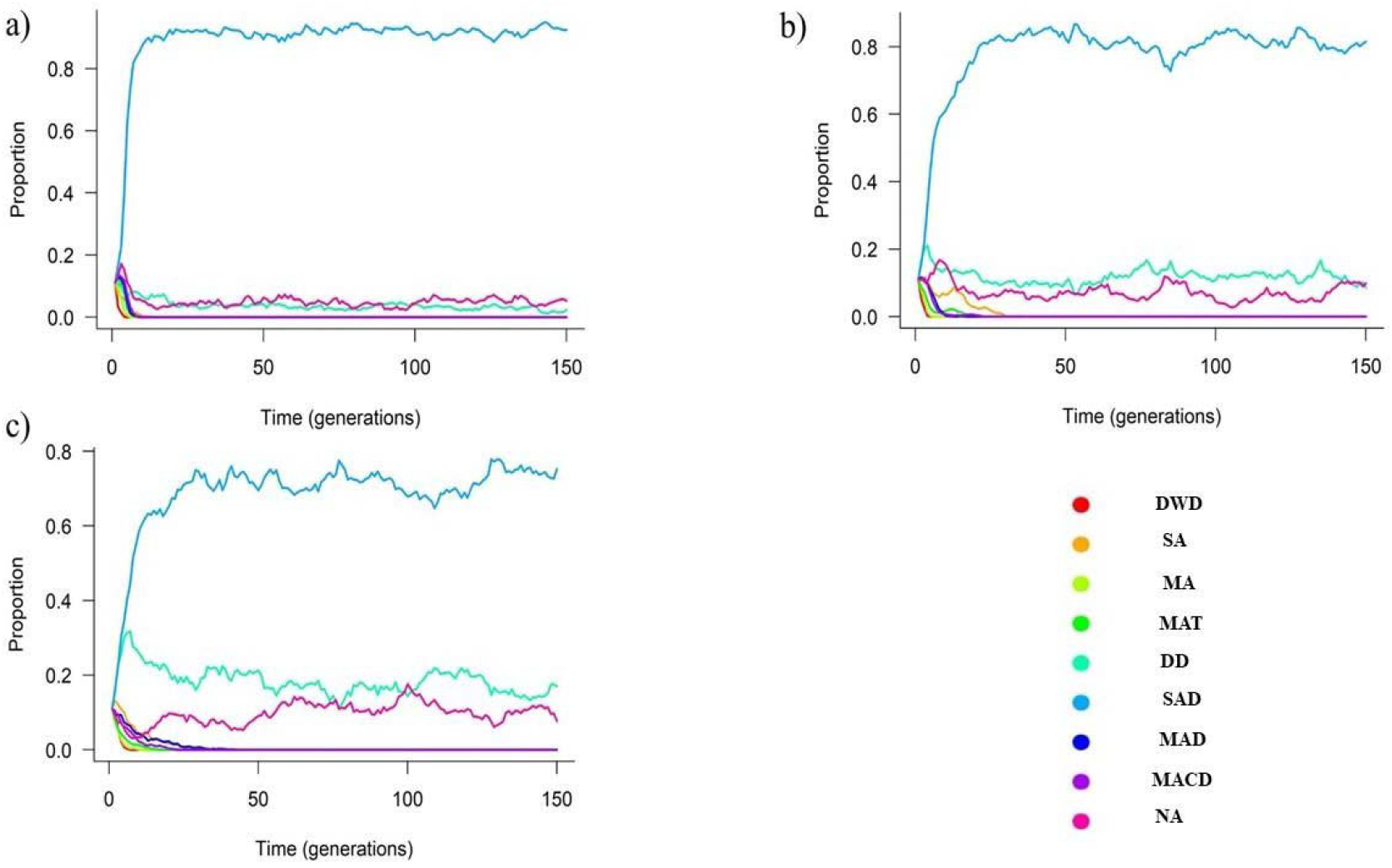
Proportion of agents with different assessment strategies during contests over resources obtained from the analysis of all generations. For all figures, the damage per round was 0.1, the probability of reproduction with the territory was 90% and without the territory was 0%. In Figure 2a, we simulated a scenario with low resource availability (N=180), Figure 2b with intermediate resource availability (N=450), and Figure 2c with high resource availability (N=900). Lines represent the proportion for each strategy. Red line: desperado without damage (DWD). Orange line: self-assessment without damage (SA). Light green line: mutual-assessment without damage (MA). Green line: mutual-assessment with cost threshold (MAT). Green tea line: desperado with damage (DD). Light blue line: self-assessment with damage (SAD). Blue line: mutual-assessment with damage (MAD). Purple line: mutual-assessment with cost threshold and damage (MACD). Pink line: non-aggressive (NA).

### **Second scenario:** high damage costs per round of contest (D=0.5) and no probability of reproduction without resource possession

For low resource availability (R=180), 79% of the agents presented self-assessment with damage strategy (Fig. 3a), 19.11% presented the non-aggressive strategy (Fig. 3a) and 1.89% presented the desperado with damage strategy (Fig. 3a). For intermediate resource availability (R=450), 64.5% of the agents presented self-assessment with damage strategy (Fig. 3b), 18.33% presented the non-aggressive strategy (Fig. 3b) and 17.17% presented the desperado with damage strategy (Fig. 3b). For high resource availability (R=900), 60.33% of the agents presented self-assessment with damage strategy (Fig. 3c), 20.28% presented the desperado with damage strategy (Fig. 3c) and 19.39% presented the non-aggressive strategy (Fig. 3c).

**Figure 3.**
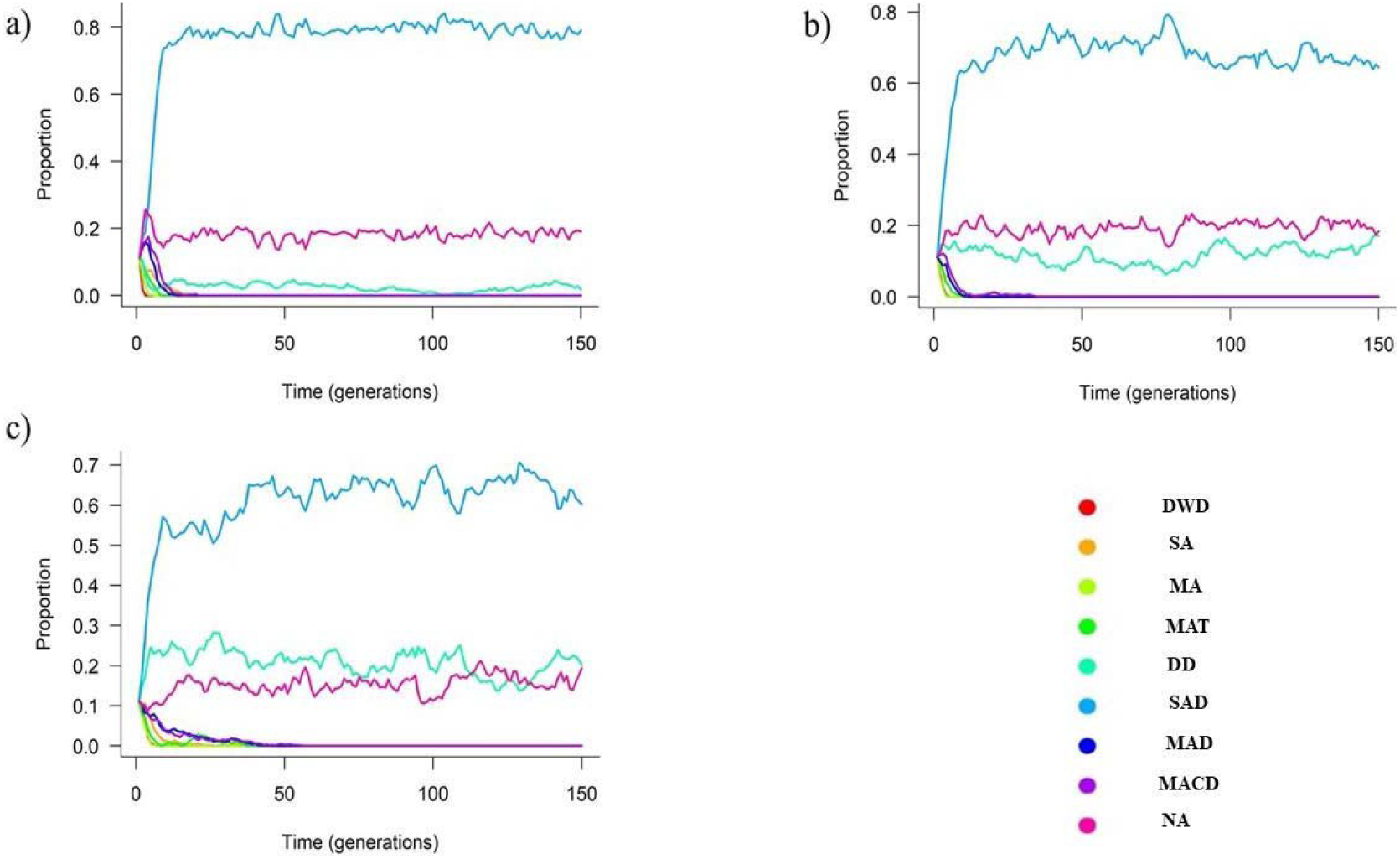
Proportion of agents with different assessment strategies during contests over resources obtained from the analysis of all generations.For all Figures the damage per round was 0.5, the probability of reproduction with the resource was 90% and without the resource was 0%. In Figure 3a, we simulated a scenario with low resource availability (N=180), Figure 3b with intermediate resource availability (N=450), and Figure 3c with high resource availability (N=900). Lines represent the proportion for each strategy. Red line: desperado without damage (DWD). Orangeline: self-assessment without damage (SA). Light green line: mutual-assessment without damage (MA). Green line: mutual-assessment with cost threshold (MAT). Green tea line: desperado with damage (DD). Light blue line: self-assessment with damage (SAD). Blue line: mutual-assessment with damage (MAD). Purple line: mutual-assessment with cost threshold and damage (MACD). Pink line: non-aggressive (NA).

### **Third scenario:** low damage costs per round of contest (D=0.1) and different probabilities of reproduction without (P=0.3) and with (P=0.9) resource possession

For low resource availability (R=180), 54.22% of the agents presented non-aggressive strategy (Fig. 4a) and 45.78% presented the self-assessment with damage strategy (Fig. 4a). For intermediate resource availability (R=450), 62.67% of the agents presented self-assessment with damage strategy and 37.33% presented the non-aggressive strategy (Fig. 4b). For high resource availability (R=900), 74.34% of the agents presented self-assessment with damage strategy, 25.33% presented the non-aggressive strategy and 0.33% presented the desperado with damage strategy (Fig. 4c).

**Figure 4.**
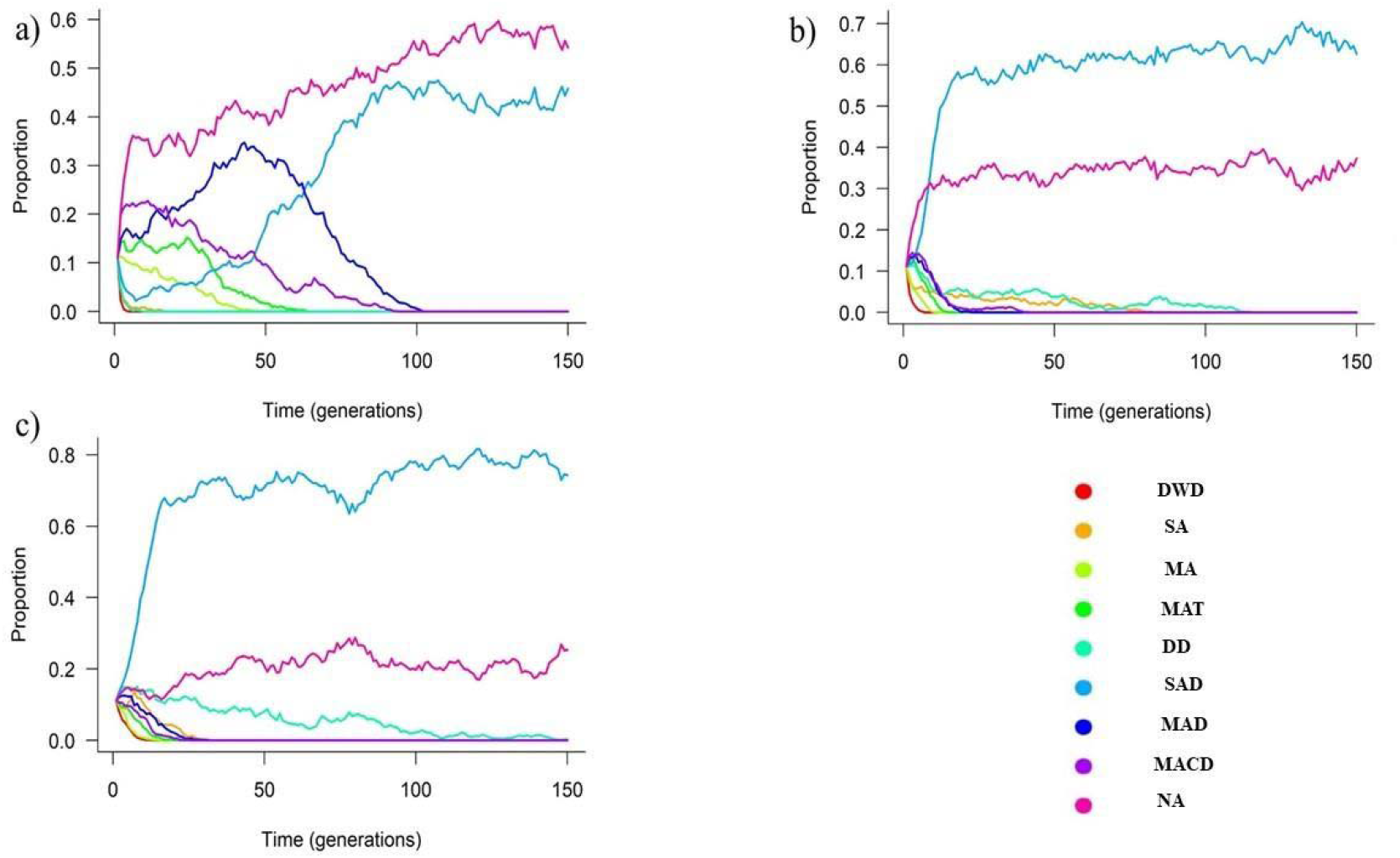
Proportion of agents with different assessment strategies during contests over resources obtained from the analysis of all generations. For all Figures, the damage per round was 0.1, the probability of reproduction with the territory was 90% and without the territory was 30%. In Figure 4a, we simulated a scenario with low resource availability (N=180), Figure 4b with intermediate resource availability (N=450), and Figure 4c with high resource availability (N=900). Lines represent the proportion for each strategy. Red line: desperado without damage (DWD). Orangeline: self-assessment without damage (SA). Light green line: mutual-assessment without damage (MA). Green line: mutual-assessment with cost threshold (MAT). Green tea line: desperado with damage (DD). Light blue line: self-assessment with damage (SAD). Blue line: mutual-assessment with damage (MAD). Purple line: mutual-assessment with cost threshold and damage (MACD). Pink line: non-aggressive (NA).

### **Fourth scenario:** high damage costs per round of contest (D=0.5) and different probability of reproduction without (P=0.3) and with (P=0.9) resource possession

For low resource availability (R=180), 88.22% of the agents presented mutual-assessment without damage strategy, 11.72% presented the non-aggressive strategy and 0.06% presented the mutual-assessment with damage strategy (Fig. 5a). For intermediate resource availability (R=450), 50.78% of the agents presented self-assessment with damage strategy, 45.83% presented the non-aggressive strategy and 3.39% presented the desperado with damage strategy (Fig. 5b). For high resource availability (R=900), 60.22% of the agents presented self-assessment with damage strategy, 30.56% presented the non-aggressive strategy and 9.22% presented the desperado with damage strategy (Fig. 5c).

**Figure 5.**
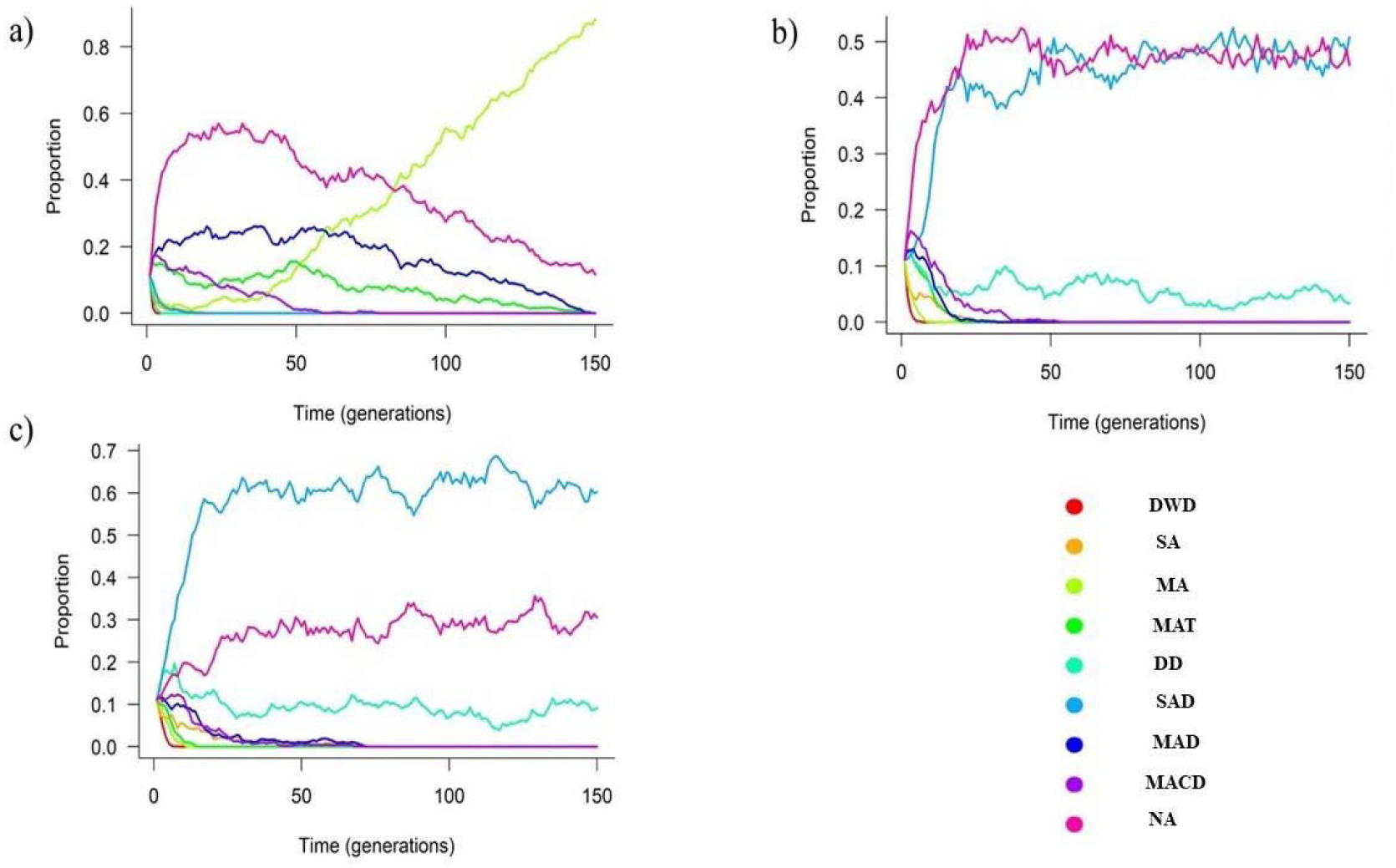
Proportion of agents with different assessment strategies during contests over resources obtained from the analysis of all generations. For all figures, the damage per round was D=0.5, the probability of reproduction with the territory was 90% and without the territory was 30%. In Figure 4a, we simulated a scenario with low resource availability (N=180), Figure 4b with intermediate resource availability (N=450), and Figure 4c with high resource availability (N=900). Lines represent the proportion for each strategy. Red line: desperado without damage (DWD). Orangeline: self-assessment without damage (SA). Light green line: mutual-assessment without damage (MA). Green line: mutual-assessment with cost threshold (MAT). Green tea line: desperado with damage (DD). Light blue line: self-assessment with damage (SAD). Blue line: mutual-assessment with damage (MAD). Purple line: mutual-assessment with cost threshold and damage (MACD). Pink line: non-aggressive (NA).

## DISCUSSION

Our ABM simulated theoretical rules proposed to understand the assessment strategies that animals may use during contests under different scenarios. We observed, for most scenarios, that agents with self-assessment with damage strategy represented the major proportion of the agents at the end of simulations. We also found that the agents using non-aggressive and desperado with damage strategies were the second and third most common strategies at most scenarios. However, they presented smaller proportion than self-assessment with damage. We found that agents using non-aggressive strategy were the major proportion of agents for only one scenario. We found that the proportion of the agents using self-assessment with damage and non-aggressive strategies presented similar proportions in only one scenario. We also found that agents using mutual-assessment without damage occurred only in one scenario. Agents with the other strategies were not represented at the end of all simulations. Taken together, our results indicated that the ability to inflict damage with a threshold of accumulated costs and a peaceful strategy may be the most common strategies under scenarios with different availability of resources and probability of reproduction with or without resources’ possession. Nevertheless, other assessment strategies can be favored depending on scenario conditions.

Self-assessment with damage strategy were prevalent in scenarios without probability of reproduction outside the resources. This pattern remained regardless of damage costs and resource availability. Then, it seems that the resource being indispensable to reproduction determines the evolution of self-assessment with damage strategy. Costlier contests may occur when the resource possession determines reproduction (Grafen 1987). However, fight using a cost threshold provided better profits than fight using a desperado strategy. Maybe, the losers using self-assessment with damage strategy can win future disputes and reproduce. On the other hand, agents using desperado with damage strategy invest all energy in one contest, decreasing the chances to win future contests and reproduce. In fact, self-assessment with damage strategy occurs in some species in which the resource possession is indispensable to reproduction. For example, males of the Wellington tree weta (*Hemideina crassidens*) fight over the possession of galleries that are indispensable to reproduction using aggressive behaviours with risk of damage (Kelly 2006). Wasps also use aggressive behaviours with risk of damage and even death during contests over hosts. It is essential for females to secure these hosts to ensure larval development, making hosts a valuable resource (Hardy et al. 2013).

Agents using the non-aggressive strategy occurred in all scenarios. Interestingly, agents using the non-aggressive strategy occurred in scenarios in which resource was essential to reproduction. However, agents using the non-aggressive strategy were prevalent only in the scenario with low resource availability, low damage and with probability to reproduce without the resource. In the scenario with intermediate availability of resources, high damage and with probability to reproduce, agents using non-aggressive and self-assessment with damage strategies were prevalent. Individuals without resources with a non-aggressive strategy can become more common in a population and may benefit from a negative frequency dependence based on the strategies followed by owners and non-owners (Kokko et al. 2006). It is possible that the agents following a non-aggressive strategy lost the resources possession at the first interactions. Nevertheless, they do not pay costs associated to damage and energy waste during the interactions. However, when agents using self-assessment with damage strategy increased their frequency in the population, it is possible that such agents had to pay higher costs due to increasing number of contests and low availability of resources. Then, the agents with the non-aggressive strategy may increase their frequencies, as they can reproduce without the resource. For example, males of some species of Odonata (e.g. *Hetaerina americana*, Raihani, Serrano-Meneses & Córdoba-Aguilar, 2008) present territorial and non-territorial tatics (Suhonen, Rantala & Honkavaara, 2008). The prevalence of males using one or another tatic in these Odonata species is partialy based on a negative frequency dependence of each tatic in the system as the observed in our results.

Mutual-assessment has been suggested as an economic way to solve contests, since individuals may flee from the contest as soon as they perceive themselves as weaker than the opponent (Arnott & Elwood 2009). Unexpectedly, agents using mutual-assessment strategies prevailed only when there was high damage, low resource availability and with probability of reproduction without the resource. The balance of high damage and low resource availability may be counterbalanced by the probability of reproduction without the resource. Then, agents using mutual-assessment strategy, due to the ability to evaluate rivals, can choose the contest that they have the best chances to win. In this scenario, it is also possible that the agents with mutual assessment ability surpassed the agents using non-aggressive strategy because they were able to reproduce either with or without the resource. However, with increasing availability of resources, agents using self-assessment with damage and the non-aggressive strategies dominated. Then, it is possible that frequency of agents with mutual assessment ability was affected by a negative density dependence (Kokko et al. 2006). Thus, although mutual-assessment was thought to be pervasive in nature (Elwood & Arnott 2012), animals using this strategy might be observed in specific situations. Therefore, it is possible that self-assessment strategies may be a general rule in animal contests (Elwood & Arnott 2012).

We conclude that alternative strategies of contest resolution occur in most scenarios and provide an evolutionary stable strategy for situations in which resource possession determines reproduction probability. One of the most prevalent alternative strategies was a strategy based on damage inflicted on other agents. On the other hand, agents using a strategy based on conflict eviction, investing all opportunities in reproduction may be also an alternative strategy to individuals that are not able to inflict damage on rivals. One can argue that these conclusions may not apply to contests in which there are no obvious ways by which rivals impose costs to each other. However, even in such contests, external costs linked to physiological stress, time wasted or even predation risk are possible and can be used as cumulative costs to rivals (Payne 1996; Payne 1998). Then, the self-assessment with damage and the non-aggressive strategies may be an evolutionary stable strategy to such scenarios. However, empirical evidence supporting cumulative assessment are related to situations in which the resource is indispensable to reproduction or to life maintenance (e.g. Pratt et al. 2003; Morrell et al. 2005; Rillich et al. 2007; Smallegange et al. 2007). It must indicate that self-assessment with damage occurs in specific conditions in natural systems. Finally, it is possible that animals may alternate strategies during the contest (Mesterton-Gibbons & Heap 2014). If the use of alternative assessment strategies during contests is common in nature, then our conclusions may be applicable to more specific conditions of natural systems.

## ACKNOWLEDGEMENTS

NSP thank CAPES for providing a PhD scholarship. I also thank to Danilo Muniz for all the help in the modelling process.

